# Colour vision is aligned with natural scene statistics at 4 months of age

**DOI:** 10.1101/2022.06.06.494927

**Authors:** Alice E. Skelton, Anna Franklin, Jenny M. Bosten

**Affiliations:** The Sussex Colour Group, University of Sussex, UK; The Sussex Vision Lab, University of Sussex, UK

## Abstract

Visual perception in adult humans is thought to be optimised to represent the statistical regularities of natural scenes (Parraga et al., 2000; Simoncelli, 2003). For example, in adults, visual sensitivity to different hues shows an asymmetry which coincides with the statistical regularities of colour in the natural world (Bosten et al., 2015). Infants are sensitive to statistical regularities in social and linguistic stimuli, but whether or not infants’ visual systems are tuned to natural scene statistics is currently unclear. We measured colour discrimination in infants to investigate whether or not the visual system can represent chromatic scene statistics in very early life. Our results reveal the earliest association between vision and natural scene statistics that has yet been found: even as young as four months of age, colour vision is aligned with the distributions of colours in natural scenes.

**Research Highlights:**

1. We find infants’ hue sensitivity is aligned with the distribution of colours in the natural world, as it is in adults
2. At just 4 months, infants’ visual systems are tailored to extract and represent the statistical regularities of the natural world
3. This points to a drive for the human brain to represent statistical regularities even at a young age.

## Introduction

Human perception is tuned in many ways to extract information from the environments that we find ourselves in. Processes to calibrate perception to the contents of natural environments occur over different time scales, both over evolutionary history (Osorio & Vorobyev, 1996; Regan et al., 2001; Changizi et al., 2006) and within the lifetime of the individual (Schefrin & Werner, 1990). Some calibrative processes undoubtedly occur during infancy. For example, perception of faces (Kelly et al., 2007), speech (Kuhl et al., 2006) and music (Trehub & Hannon, 2006) is refined during infancy to show biases for the stimuli encountered in infants’ environments. In mature perceptual systems there is tuning to the statistical regularities of visual scenes: visual channels are thought to be tuned to extract the regularities of spatial frequency (Knill et al., 1990; Tadmor & Tolhurst, 1994; Hansen & Hess, 2006), orientation (Hansen & Essock, 2004), and attributes of colour (Long et al., 2006; McDermott & Webster, 2012; Smet et al., 2016; Webster, 2015a; Webster, 2015b; Welbourne et al., 2015) in natural scenes. Relatively little is known about the developmental trajectories of visual representations of the statistical regularities of natural scenes. Ellemberg et al. (2012) have found that biases in perception according to the spatial frequency structure of natural scenes are present in 10 year-olds but undeveloped in 6 and 8 year-olds. The only existing work on infants has found that at 9-10 months, but not at 3-6 months, infants’ looking preferences and their Event Related Potential components (P100 and P400) appear to distinguish between natural textures and one type of synthetic manipulation of those textures (Balas & Conlin, 2014; Balas & Woods, 2014). One interpretation of this work is that infants tune to natural texture statistics by 9 months. However, the lack of a consistent effect of another type of natural texture manipulation (contrast polarity), and the potential role of other low-level visual differences independent of naturalness suggests that further work is needed to clarify the nature of these effects.

Colour vision is a good example of a perceptual system that is tuned at several different levels to extract information from natural scenes. Human colour vision relies on three classes of retinal receptors, long- (L), medium- (M) and short- (S) wavelength sensitive cones. The particular sensitivities of the human cones are thought to have evolved to represent colour information relevant to the diets or skin colours of our primate ancestors (Osorio & Vorobyev, 1996; Regan et al., 2001; Changizi et al., 2006). Postreceptorally, colour is processed predominantly in two chromatically opponent channels, which themselves seem to be tuned to minimise the redundancy of information acquired about natural scenes (Ruderman et al., 1998). There has been recent interest in a third level of alignment with natural scene statistics: colour discrimination is poorest along a blue-yellow colour axis (Honjyo & Nonaka, 1970; Krauskopf & Karl, 1992; Pearce et al., 2014; Bosten et al., 2015; Álvaro et al., 2017), which is intermediate in colour spaces between the colour axes along which the two chromatically opponent retinogeniculate colour mechanisms are tuned (the vertical and horizontal axes in Figure 1a). It is the blue-yellow axis that contains most variance in natural scenes (Webster & Mollon, 1997; Bosten et al., 2015), and it has been proposed that a trade-off between the range of colours that a colour mechanism must represent and the accuracy with which it can represent them may explain why discrimination is worst along this blue-yellow axis (von der Twer & MacLeod, 2001; MacLeod & von der Twer, 2003; Bosten et al., 2015).

**Figure 1.**
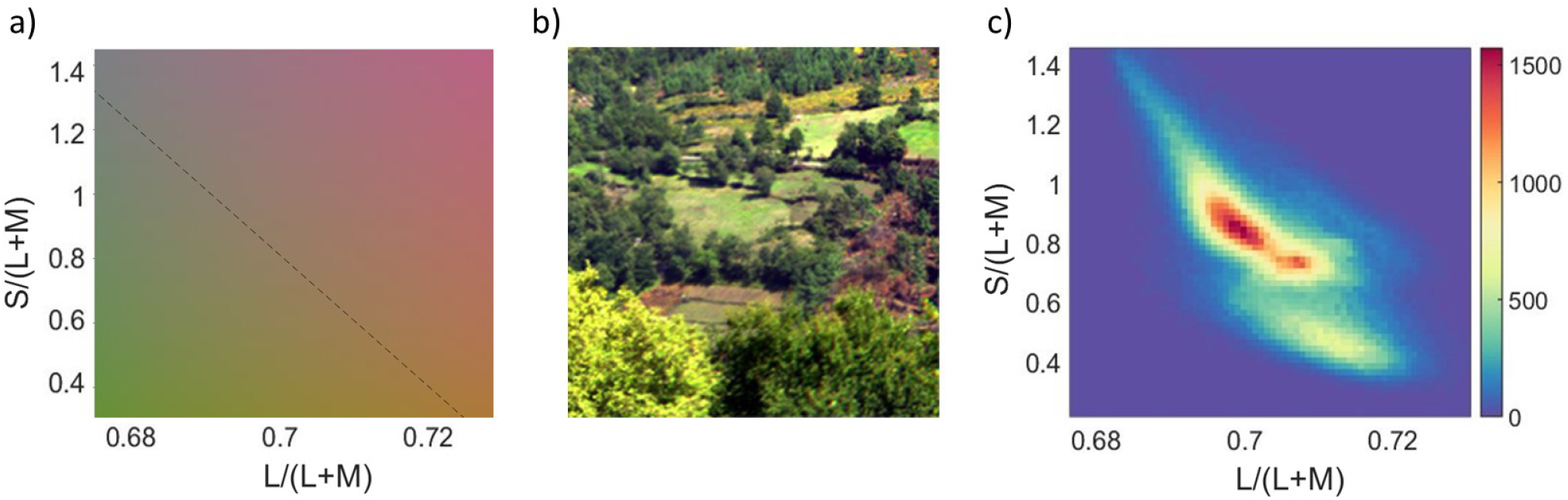
(a) Our version of the MacLeod-Boynton (1979) chromaticity diagram, constructed using the Stockman, MacLeod and Johnson (1993) cone fundamentals, representing the signals carried by the chromatically opponent retinogeniculate colour mechanisms, L/(L+M) and S/(L+M). The blue-yellow axis, indicated by the dashed line, is defined as the line connecting unique blue at 476 nm and unique yellow at 576 nm (Mollon, 2006) and is intermediate to the cardinal axes. (b) An example of a hyperspectrally imaged natural scene (Nascimento et al., 2002), and (c) A histogram showing the distribution of chromaticities of the natural scene in (b) plotted in our version of the MacLeod-Boynton chromaticity diagram. The largest variance in chromaticities is along the negative diagonal.

In the present study we investigated the development of representations of the distribution of colours in natural scenes by measuring colour discrimination in infants using a novel task. We chose infants aged 4-6 months, because it is thought that the S-cone reliant colour-opponent mechanism develops relatively late at 2-3 months (Bornstein, 1975; Brown & Teller, 1989; Knoblauch et al., 2001; Suttle et al., 2002), and it is only after this that infants have trichromatic colour vision. For 60 infants and 19 adults we measured eye movements to chromatic targets varying across 8 levels of saturation in 8 hue directions.

## Methods

### Participants

Sixty-four infants (25 female, age range 4.01–6.80 months, mean age 5.18 months, SD 0.76 months) took part in the experiment. Four infants were excluded from the study due to fussiness preventing them from proceeding past the initial calibration phase. All participants were born at 40±2 weeks, had a minimum birth weight of 2500 g, had no known visual deficits, no history of epilepsy, and reported no family history of colour vision deficiency.

Nineteen adult participants took part in the study (13 female, mean age 26.42 years). Participants had normal colour vision as assessed by the Ishihara Plates test (Ishihara, 1917) and the Lanthony Tritan Album (Lanthony, 1987).

Ethical approval for the study was granted by the University of Sussex Science and Technology Cross-Schools Research Ethics Committee (ER/AF247/17), and the European Research Council Executive Agency.

### Stimuli and apparatus

Target stimuli were coloured circles that subtended a visual angle of 6.9° on a surround metameric with equal energy white that subtended a visual angle of 41°. The chromatic difference between target and surround was isoluminant for the standard observer (Stockman et al., 1993). In order to prevent individual participants detecting targets via residual luminance differences with the surround, low-pass luminance noise was added over the whole display. The mean luminance of both target and surround was 28 cdm^-2^, and luminance noise was linearly distributed to range between ±10% of the mean luminance. The chromaticities of the target stimuli varied in saturation along 8 axes in a version the MacLeod-Boynton (1979) chromaticity diagram based on the Stockman, MacLeod and Johnson (1993) cone fundamentals (Figures 2a and b). The centre of our stimulus space was metameric with equal energy white (the background chromaticity), and there were 8 points per axis (64 points in total) increasing in saturation away from the centre. Four of the axes were along the cardinal directions in the MacLeod-Boynton chromaticity diagram; two axes were in the blue-yellow direction (the blue-yellow line was defined from 476 to 576 nm; Mollon, 2006), and two axes (lying close to the positive diagonal) were a reflection of the blue-yellow line through the vertical axis of the MacLeod-Boynton chromaticity diagram.

**Figure 2.**
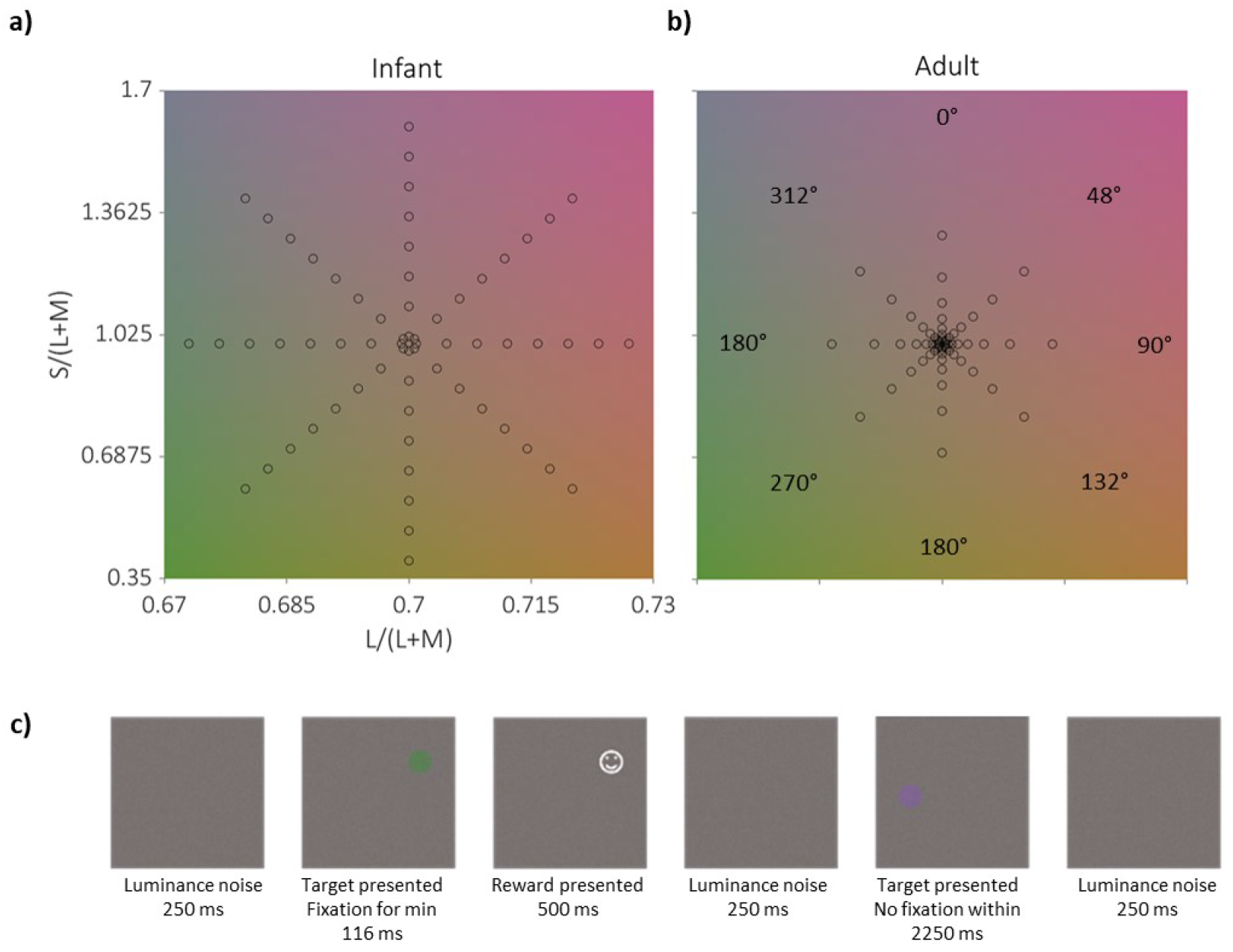
Target chromaticities for (a) infants and (b) adults along the 8 axes plotted in the MacLeod-Boynton chromaticity diagram. The hue angles of the 8 axes are indicated in (b). Note that the axes near the positive and negative diagonals were defined as the blue-yellow axis translated to the white point and its reflection, and not as the mid-points of the cardinal axes. Towards the centre of the figure, colours are desaturated, and increase in saturation radially along the axes. (c) Trial procedure. Participants were presented with a blank screen of luminance noise for 250 ms before a target was presented. If the target was fixated for a minimum of 116 ms, the target was replaced by a smiley face, a sound reward was presented, and a hit was recorded for the trial. After another blank display of luminance noise, the next target was displayed. If the target was not fixated within 2250 ms, no reward was presented, and the next trial began.

For adults, stimuli were spaced logarithmically along these axes, to focus the collection of data closest to where adult thresholds were expected to be (Figure 2b). For infants, stimuli were sampled in equal steps along each axis (Figure 2a). Due to infants’ higher colour discrimination thresholds (Knoblauch et al., 2001) stimuli were sampled from a wider range of chromaticities than for adults. The maximum target saturations for infants were limited by the display’s colour gamut.

Stimuli were presented on a Diamond Pro 2070SB CRT Monitor (Mitsubishi, Tokyo, Japan) via a PC-driven ViSaGe MKII Stimulus Generator (Cambridge Research Systems Ltd, Rochester, UK). Eye-movements were recorded via an Eyelink 1000 system (SR Research, Ontario, Canada), and the experimental programme was run in MATLAB (R2016a, The MathWorks Inc., Natick, MA). The display was gamma-corrected using a LS-100 luminance meter (Konica-Minolta, Tokyo, Japan), and the colour calibration was achieved using a Spectrascan PR655 spectroradiometer (Photo Research Inc., Chatsworth, CA).

### Procedure

Infant participants were seated in a car seat mounted on a chair 60 cm from the display. They watched a cartoon on the display during camera set up, and then completed a 9-point spatial calibration. If necessary (e.g., if a participant moved substantially), the spatial calibration was repeated during testing.

Before each trial, participants were presented with a blank screen of luminance noise for 250 ms. Targets appeared within 6.9° of the participant’s current fixation point on the display. If the participant fixated the target (or a 1 ° area around the target) for at least 116 ms, this was recorded as a successful fixation (a ‘hit’), the target was replaced with an image of a smiley face, and a short musical tune was played. If the target was not fixated within 2250 ms, no auditory or visual reward was played, and a miss was recorded (Figure 2c).

If an infant participant’s gaze left the screen, a looming black and white spiral accompanied by a loud noise was presented to refocus visual attention to the screen. Once the participant was centrally fixated, the experimenter pressed a key to allow trials to start again. In order to encourage maximum engagement with the task for infants, informed by pilot testing (N=4), a maximally saturated stimulus was presented on 3 out of every 10 trials. Stimuli were presented in blocks of 64, and testing was continued until the infant became disengaged with the task.

For adult participants, trials were presented in 10 blocks of 64 trials (640 trials in total), and the test duration was approximately 30 minutes. During each block of trials, each chromaticity was presented once, in a random order, so each chromaticity was presented 10 times to each participant. Participants were told that their task was to continually search for and look at targets on the screen.

### Analysis

For each hue axis, a local linear *Modelfree* algorithm (Zychaluk & Foster, 2009) was used to fit a psychometric function to the rate of detection plotted against target saturation (an example psychometric function is shown in Figure 3a). A colour discrimination threshold can be defined at a certain performance criterion (alpha) along the psychometric function. Through thresholds for each of the eight colours we fit colour discrimination ellipses, where the distance from the centre of the ellipse in each hue direction is the discrimination threshold in our version of the MacLeod-Boynton (Macleod & Boynton, 1979) chromaticity diagram. We chose the MacLeod-Boynton chromaticity diagram as a colour space because it represents along its cardinal axes activity in the two dominant post-receptoral retinogeniculate colour mechanisms (Webster & Mollon, 1991). If colour discrimination were simply limited by the signal to noise ratio available in the low-level retinogeniculate colour mechanisms, colour discrimination ellipses plotted in the MacLeod-Boynton (1979) chromaticity diagram would be oriented either horizontally or vertically. An off-axis bias in colour discrimination would reveal the action of non-cardinal colour mechanisms such as those that may be tuned to the blue-yellow statistics of natural scenes (Bosten et al., 2015).

**Figure 3.**
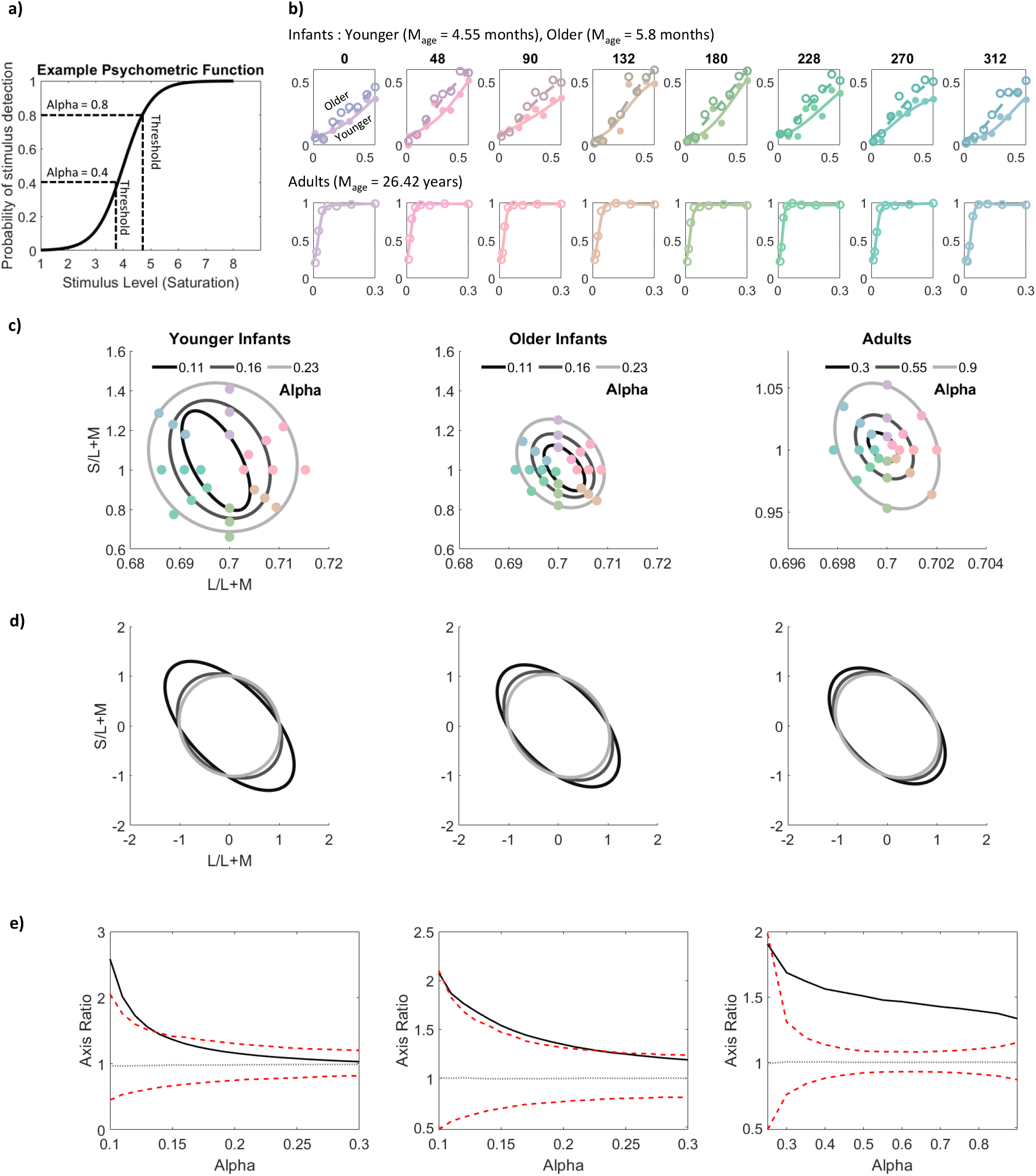
(a) Example psychometric function. (b) psychometric functions for infants (upper panels) and adults (lower panels) for the 8 colour axes tested, all showing the probability of target detection as a function of stimulus intensity (saturation). The example psychometric function demonstrates how different alpha criteria result in different thresholds along the same psychometric function. For the infant and adult data, each individual graph shows group average psychometric functions for a specific hue axis, which is a radial line in the MacLeod-Boynton chromaticity diagram (Figure 2a and b), at an angle specified in the bottom left of each figure (For correspondence between angles and axis, see Figure 1b). (b) For each of a set of alphas, 8 colour discrimination thresholds were extracted (one for each hue), and then an ellipse was fit to each set of 8 thresholds plotted in the unnormalised MacLeod-Boynton chromaticity diagram. Separate discrimination ellipses were plotted for the 3 groups: two infant groups (younger and older than the infant sample median age), and adults; note that the axis limits are different between infant and adult panels because of the large group difference in mean thresholds. The figures show ellipses for 3 example alphas. (c) In order to quantify the extent of the blue-yellow bias of each colour discrimination ellipse, an ‘axis ratio’ was calculated in the normalised version of the MacLeod-Boynton chromaticity diagram, where the ellipse is scaled so that it’s x- and y-intercepts are at -1 and 1 when it is centred at (0,0). The axis ratio in this space is the ratio of the length of the negative diagonal of the ellipse to the length of the positive diagonal. In (b) and (c), ellipses fit to thresholds extracted at lower alphas are shown lighter in colour, and those from higher alphas are shown darker in colour. (d) Axis ratios are plotted as a function of alpha for each of the three groups (solid black lines). The range of 95% of permuted functions are shown by the red dashed lines. The median permutation is shown by the dotted grey line (see text for details of the permutation analysis).

From the fitted ellipses we quantified the asymmetry in colour discrimination by taking an *axis ratio*, calculated after ‘normalising’ the ellipses so that, when centred at (0,0), their x- and y-intercepts were 1 and -1 (Bosten et al., 2015). For normalised ellipses, the axis ratio is the ratio of the length of the ellipse along the negative diagonal to the length along the positive diagonal. Normalised ellipses are circular and have an axis ratio of 1 if the un-normalised discrimination ellipses are oriented horizontally or vertically in the MacLeod-Boynton (1979) chromaticity diagram. If the axis ratio is greater than 1, then colour discrimination is relatively poor in a blue-yellow colour direction, aligned with the axis of maximum variability of colours in natural scenes. If the axis ratio is smaller than 1, then colour discrimination is relatively poor in an orthogonal roughly red-green direction.

## Results

For our primary analysis we fit colour discrimination ellipses for three groups, two infant groups split above and below the median age (younger infants: N = 30, M_*age*_ = 4.55 months, SD = 0.27, and older infants: N = 30, M_*age*_ = 5.82 months, SD = 0.51), and adults. Due to missing data (infant participants were not expected to complete all 640 possible trials) it is not possible to calculate a threshold for each infant for every hue angle, however see SI Section 2 for results extracted on an individual level.

For each of the three age groups and for each hue, we achieved very clear group psychometric functions (Figure 3b). One obvious difference between the groups is in the positions of the asymptotes of the psychometric functions. For adults, performance quickly asymptotes close to 100% detection. For younger infants the asymptotes are at about 30-40% and for older infants they are at about 50-60%. This large difference means that any particular performance criterion (alpha) would fall on very different parts of the psychometric function for the different groups. This prompted us to extract thresholds at different alphas (threshold criteria), and to fit different discrimination ellipses to each set of thresholds at different alphas (Jones et al., 2015; Trehub et al., 1980). Figure 2(c) shows example colour discrimination ellipses for each group defined at three different alphas, and the same results are plotted in Figure 2(d) in the normalised version of the MacLeod-Boynton chromaticity diagram in which we extract axis ratios. All the colour discrimination ellipses are elongated along the negative diagonal, particularly those at lower alphas (plotted in darker grey in the figure), and therefore have axis ratios greater than 1. We also note evidence for improving colour discrimination with age in agreement with earlier literature (Knoblauch et al., 2001): discrimination ellipses for younger infants are larger than for older infants (see SI Section 3 for an analysis of the relationship between colour discrimination thresholds and age).

For each group Figure 3(e) shows axis ratio plotted as a function of alpha. For all three age groups, axis ratio is larger at low alphas and then decreases as a function of alpha. Our choice of *Modelfree* local linear fits for our psychometric functions has not been critical to our results: similar results are obtained when parametric cumulative Gaussians are fit (see Figure SI.1 for results using *Modelfree* local linear fits and results using cumulative Gaussian fits shown on the same axes). Also plotted in the figure is the results of a permutation analysis where the analysis was re-run 1000 times but with the connection between hue angles and responses disrupted by permutation. The dashed red lines indicate the range of axis ratios as a function of alpha in 95% of permutations. For all three groups the observed axis ratios are larger than 95% of permuted axis ratios at least at some alphas, showing that colour discrimination is selectively poor in the blue-yellow colour direction, as would be expected if colour vision is aligned with the statistics of natural scenes. For infants, and particularly for younger infants, axis ratios are large at low alphas, but as alpha increases they tend towards 1. Adults also show a decrease in axis ratio with alpha, but maintain an axis ratio larger than one at all alphas. We conducted similar analyses at the level of individual participants for infants as one group and adults as a second group (for details see SI Section 2).

## Discussion

Our results show that like adults, infants as young as 4 months of age show a bias in colour discrimination in a direction predicted from the distributions of colours in natural scenes. Though there is a significant improvement in colour discrimination with age (see SI Section 3), the discrimination ellipses show no evidence for development in the degree of bias (Figure 3b). Our results are compatible with two schemes: either the blue-yellow bias in colour discrimination is innate, perhaps calibrated to natural scene statistics neurogenetically, or it develops very early by calibration to the statistics of natural scenes, between the ages of 2-3 months (when trichromacy first develops) and 4 months (the age of the youngest infants in our sample).

The dependence of axis ratio on alpha that we observe for infants implies that the psychometric functions describing performance as a function of saturation have both higher mean saturations and steeper slopes for blue and yellow targets than for red and green targets. This does not fit the usual expectation, where the noise that limits detection tends to increase the mean (determined by signal:noise ratio) and reduce the slope (Arazi et al., 2017; Legge et al., 1987). It is possible that the greater amount of noise in threshold estimates at low alphas has influenced the axis ratios we observe (the range of axis ratios calculated from permuted data is greater at lower alphas; Figure 3e). However, it may also be the case that non-visual processes contribute differently to target detection at different alphas and may thus differently affect axis ratio depending on alpha. Specifically, it is generally assumed that the slope of the psychometric function is determined by visual signals and sensory noise, while attentional ‘lapses’ affect only its upper asymptote (Swanson & Birch, 1992; Wichmann & Hill, 2001). However, especially for infants, this may not be a safe assumption. Instead, there may be an interaction between visual and non-visual contributions to psychophysical decisions so that both the positively sloping region and the upper asymptote are influenced by non-visual factors such as attention and motivation. The non-visual factors may have their greatest influence at high signal strengths. For example, in the current study, an infant may be more motivated (e.g., by attentional salience) to make an eye movement towards a coloured stimulus with greater saturation than one with lower saturation. Thus, the lower reaches of the psychometric function at lower stimulus strengths may be more singularly determined by visual sensitivity than the upper reaches, where the presence or absence of attention and motivation may have a relatively greater influence on the probability of detection. Axis ratio may be largest at low stimulus strengths because this is where visual sensitivity (tuned to natural scene statistics) is having its purest influence on detection performance. Another indication of the influence of non-visual factors on the psychometric function is the fact that for infants they did not asymptote within the range of contrasts tested (Figure 3a), but were still increasing even for colours at edge of the display’s colour gamut. It seems likely that infants are visually sensitive to these stimuli but that their performance is limited by attention or task engagement.

Our finding that bias in colour discrimination varies with threshold criterion (alpha) for all three age groups has implications for psychophysics. The choice of alpha in psychophysical analysis is often arbitrary, but there are conventions that are applied. For example, alpha may be defined half way between the guess rate and the asymptote of the psychometric function (Watson & Pelli, 1983). Our finding implies that the choice of alpha may have an impact on the conclusions of psychophysical studies, and that actually a lower alpha than is typically chosen may better isolate the sensory processes (at least in infants) that psychophysicists are interested in.

The relationship between axis ratio and alpha that we have observed would not have been apparent had we not measured full psychometric functions. Typically, in infant studies, discrimination performance is measured using a variant of the ‘preferential looking’ technique, where gaze durations in the presence and absence of stimulus targets are compared statistically. This often results in one number to decide whether a given target is above or below threshold (seen versus not seen). Discrimination thresholds have been measured using preferential looking for infants of different ages (Knoblauch et al., 2001) but owing to the time-consuming nature of preferential looking, are available only for a limited number of colour axes (see SI Section 3). Our method of using eye movements (see also Jones et al., 2014) is more efficient, allowing measurement of performance at a greater number of points and full psychometric functions, as is good practice for psychophysics in adults. Our relatively efficient measure enables infants’ colour discrimination ellipses to be revealed for the first time.

Our results provide the first evidence that post-receptoral vision is aligned to natural scene statistics in early infancy. A major finding in colour vision science is that the L-, M- and S- cones are spectrally positioned to extract relevant colour information from the natural world (Osorio & Vorobyev, 1996; Regan et al., 2001; Changizi et al., 2006). If the alignment that we see in infants is achieved by neurogenetic calibration, then there must be genetic optimisation post-receptorally as well. If the aligning process is ontogenetic, then the infant brain must be able to extract and calibrate to statistical information within a matter of weeks. This revises previous research that has demonstrated statistical optimisation of perception only later in infancy for social or linguistic stimuli (Kuhl et al., 2006; Kelly et al. 2007). To distinguish these theoretically important accounts, future research will need to apply our methods to genetically similar infants living in radically different visual environments.

## Acknowledgements

We thank the infant participants and their parents, Kenneth Knoblauch, Daniel Osorio, and Donald MacLeod for helpful discussions. This research was funded by European Research Council grant 772193 COLOURMIND awarded to Anna Franklin.

