## Supplementary material for "Colour vision is aligned with natural scene statistics at 4 months of age": SI

### 1. **Data and analysis**

#### *1.1 Infant Data*

For analysis at a group level, infants were split by median age into two groups (younger and older than the median age). Details of the mean number of trials completed by each infant group and the mean proportion of ‘seen’ targets for each group are provided in Table SI.1.

**Table SI. 1. The mean number of trials and proportion of targets seen (‘hits’) across all trials by all infants, younger, and older infants.**

| Group | Mean age months (SD) | Mean number of trials (SD) | Mean proportion of ‘hit’ trials (SD) |
| --- | --- | --- | --- |
| All infants | 5.18 (0.76) | 118.25 (55.89) | 0.24 (0.15) |
| Younger infants | 4.55 (0.27) | 106.23 (48.45) | 0.20 (0.13) |
| Older infants | 5.82 (0.51) | 130.27 (60.90) | 0.29 (0.15) |

#### *1.2 Fitting psychometric functions to infant and adult response data*

For each hue axis, a psychometric function was fit to the proportion of target hits at each saturation level using a *Modelfree* local linear fit (Zychaluk & Foster, 2009) with a logit link function. The algorithm was provided with estimated lapse and guess rates. The lapse rate was 0.0242 for adults, and 0.3 for infants, estimated as the proportion of maximally saturated targets missed (over all hue angles). The guess rate was 0.0412 for infants and 0.197 for adults, estimated from the proportion of hits for minimally saturated targets (over all hue angles). Our choice of *Modelfree* local linear fits to fit psychometric functions did not substantially influence our results. We obtained similar results when using a parametric cumulative Gaussian fit, available as part of the same *Modelfree* toolbox (Zychaluk & Foster, 2009). Figure SI.1. shows a comparison of the relationship between axis ratio and alpha obtained for local linear fits and that obtained for cumulative Gaussian parametric fits.

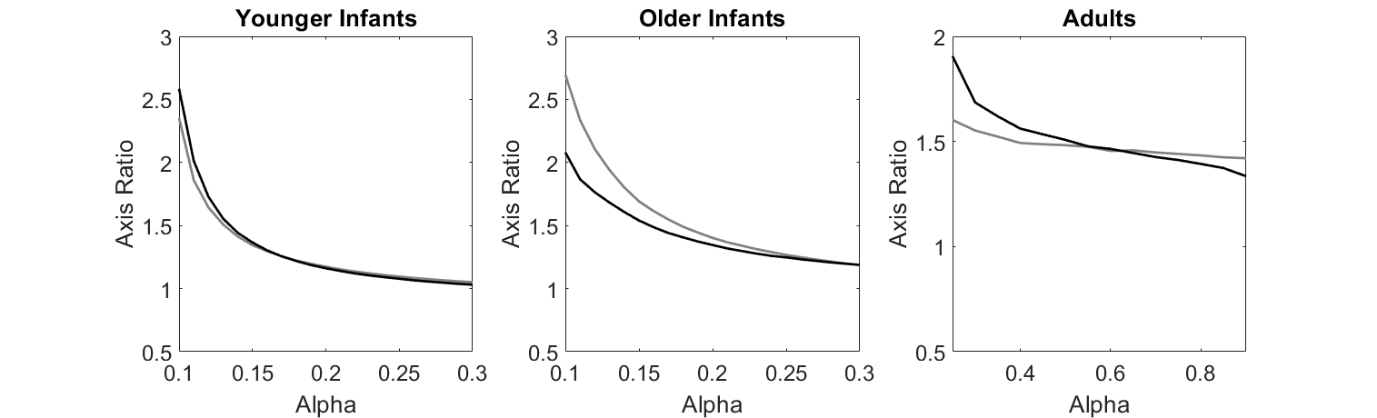

**Figure SI.1. Comparison of the observed relationship between alpha and axis ratio for *Modelfree* local linear fits (black lines) and for cumulative Gaussian parametric fits (grey lines).**

Through thresholds for the eight colour axes, at each chosen alpha we fit a colour discrimination ellipse, where the distance from the centre of the ellipse in each hue direction is the discrimination threshold in our version of the MacLeod-Boynton (1979) chromaticity diagram based on the Stockman, MacLeod and Johnson (1993) cone fundamentals.

### **2. Individual participant analysis**

#### *2.1 Individual infant analysis*

Infant participants were not expected to complete all 640 possible trials. This ‘missing data’ means it is not possible to calculate a threshold for each infant for every hue angle. In order to extract axis ratios for individual infants, data from the two hue angles along the blue-yellow axis (132^o^ and 312^o^) were collapsed together, to maximise the data available for that colour direction, and then a psychometric function was fit. The same was also done for the two hue angles of the positive diagonal (48^o^ and 228^o^). As a result there were a greater number of data points for each saturation along the positive and negative diagonals than when fitting to individual hue angles, improving the ability to fit psychometric functions to an individual infant’s data. The same *Modelfree* local linear fitting algorithm was used, but guess rates and lapse rates were free parameters as there were not enough data to estimate these for individual infants.

Nine infants were excluded from this analysis as despite collapsing along the axes, they did not have sufficient points to fit a psychometric function, either for the primary analysis or for further analysis where the connection between hue angles and response was permuted. We defined axis ratio as the ratio of the threshold along the negative diagonal to the threshold along the positive diagonal. As for the group-level analyses, different axis ratios were extracted for different alphas. Axis ratios which were particularly large (greater than 10) or small (smaller than 0.1) were excluded from further analysis. Example psychometric functions for an individual infant, and an example of the relationship between axis ratio and alpha for an individual infant are shown in Figure SI.2.

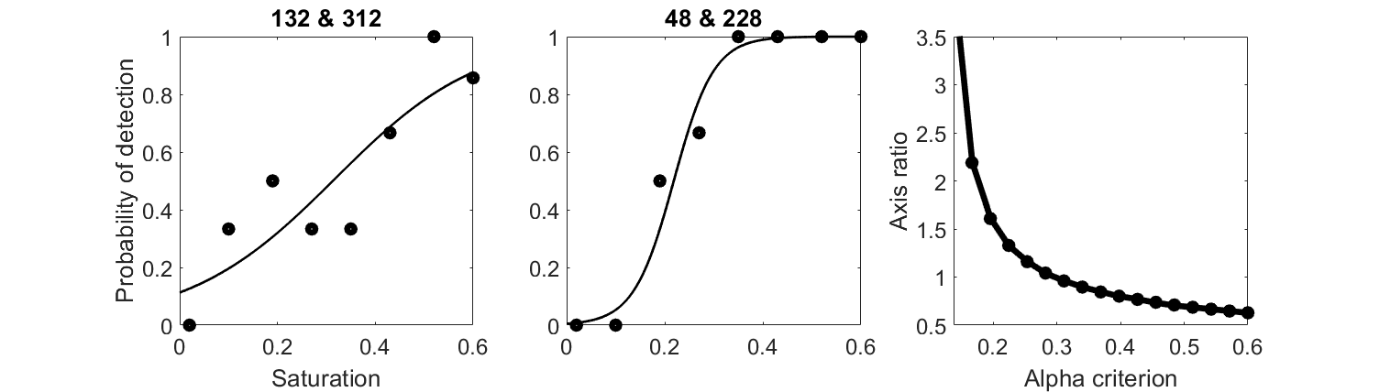

**Figure SI.2. Example psychometric functions for an individual infant (aged 5.22 months)) along the negative (48-228) diagonal (left panel) and positive (132-312) diagonal (centre panel) of the MacLeod Boynton chromaticity diagram, and the resulting axis ratios as a function of alpha (right panel).**

We also conducted a permutation analysis similar to that done for the group results (main text). On each of 1000 permutations the relationship between hue angle and response was permuted but the other stages of the analysis were maintained. Group mean axis ratios as a function of alpha are shown in Figure SI.3 (left panel), along with the range of 95% permuted functions. The observed average function from 51 individual infants does not fall outside the range of 95% permuted functions. Although there is a trend for axis ratios >1 and for a reduction in axis ratio with alpha, there is too much noise in data from individual infants to draw statistical conclusions from the individual results.

*2.2 Individual adult analysis*

We conducted an analysis of results from individual adults in the same way as for individual infants. For the local linear fits lapse and guess rates were not free parameters but estimated for individual adults because there were sufficient data to allow this. For each adult, guess rate was estimated as the proportion correct for the targets of minimum saturation (over all hue angles) and laspse rates were estimated as the proportion of misses for targets of maximum saturation (over all hue angles).

The mean relationship between alpha and axis ratio from the results of individual adults is shown in Figure SI.3 (right panel). The observed results fall outside 95% of permuted results, indicating that axis ratios for individual adults tend to be significantly greater than 1, and that colour discrimination is worse along roughly the blue-yellow axis (negative diagonal of the MacLeod-Boynton chromaticity diagram) than along the red-green axis (positive diagonal).

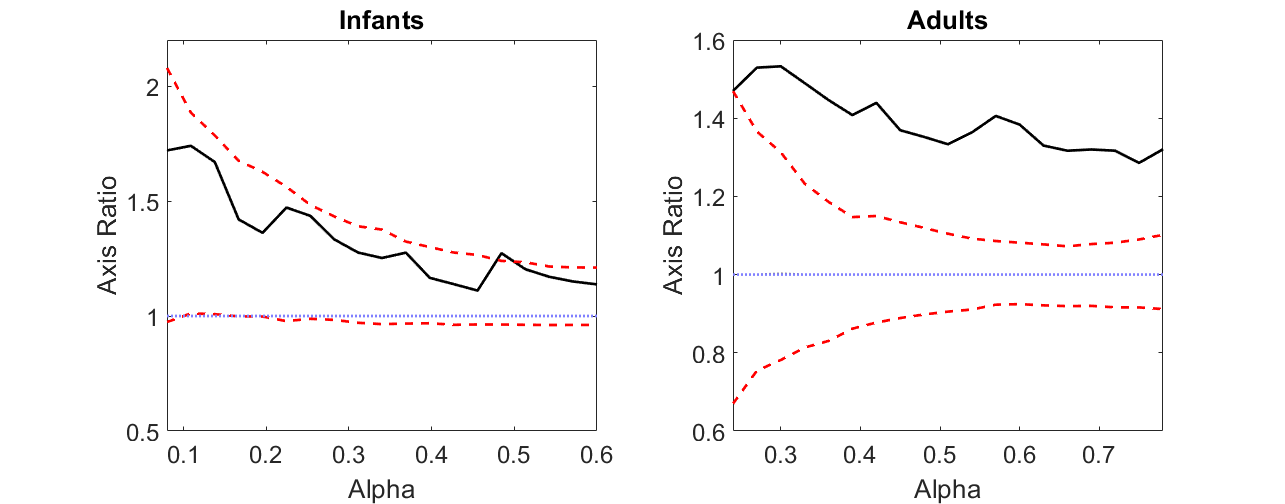

**Figure SI.3. Results of an analysis of the data from individual infants (left panel) and individual adults (right panel). In both panels, the black solid line shows the average relationship between alpha and axis ratio, and the red dashed lines show the range of 95% of permuted results.**

**3. Development of colour sensitivity**

Previous studies measuring chromatic discrimination in infants are sparse (Knoblauch et al., 2001; Teller et al., 1998). One systematically measures chromatic discrimination along three colour axes (Knoblauch et al., 2001). The three axes alone are not able to provide information about the discrimination ellipse of infant participants. Our results when plotted in the same chromaticity diagram were consistent with the thresholds found in this previous study. The method of the current study is able to capture chromatic thresholds from a greater number of directions in colour space than have previously been tested, and can be used to create full colour discrimination ellipses. This new method allows measurements to be taken with relative ease and efficiency, and allows both individual and group-level analyses of performance.

Saturation thresholds are, of course, lower at lower alphas than at higher alphas (Figure SI.4). Psychophysical methods which use a single alpha criterion to report infant discrimination thresholds may be not fully capture infant perception. The youngest infants’ performance only reached about 0.35 even for maximally saturated targets at the edge of the colour gamut (see Figure 2(b) in the main text for a comparison of psychometric functions for older and younger infants). This highlights that in interpreting the development of saturation thresholds across infancy, the co-development of attentional factors which can influence infant behaviour should also be considered. It may be that the development of these two factors differs. If the factors determining thresholds at are identical at each alpha criterion then the rate at which target detection improves with age should remain the same at all alpha criterion levels – performance would be limited by the same processes regardless of where along the psychometric function the threshold was taken. This would be represented by identical slopes in each sub panel below (Figure SI.3). However, if visual factors dominate at low alphas in determining detection thresholds, and attentional factors dominate at high alphas, then different development of these two factors might show in Figure SI.4 as different slopes in different sub-plots. There is some suggestion of this in our results, for example, the slope at alpha = 0.35 is greater than at lower alphas (e.g., 0.08).

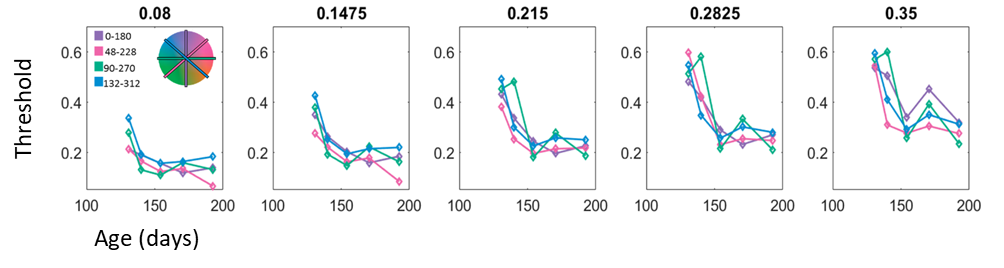

**Figure. SI.4. Development of colour discrimination thresholds along 4 major axes in the MacLeod Boynton chromaticity diagram. The key in the left panel provides details of the hue angles included for each axis, and the inset colour space shows their location in the MacLeod Boynton chromaticity diagram. Each panel represents thresholds taken at a specific alpha, identified above each panel. The development of saturation thresholds occurs at the same rate for all four axes in the MacLeod Boynton chromaticity diagram (as indicated by the overlapping lines at all alphas). Although saturation thresholds progress at the same rate across different axes, the difference in slope between panels (i.e. at different alpha criteria) suggests that there may be variability in the developmental trajectories of different factors which affect target detection, such as attention and visual sensitivity.**
